# MARRI: Mechanism-Aware Learning of RNA–RNA Interactions with Foundation Models

**DOI:** 10.64898/2026.11.17.767455

**Authors:** Bo Wang, Haozheng Luo, Zhaohan Meng, Gaofei Chen, Xiaoyu Wang, Yan Chen, Jiangning Song, Joseph Loscalzo, Hao Xu

## Abstract

RNA–RNA interactions (RRIs) are central to post-transcriptional regulation, yet their prediction requires accounting for structural accessibility and intermolecular base pairing beyond sequence complementarity. Pretrained RNA foundation models provide representations of individual RNAs, but translating these representations into interaction predictions requires mechanistic knowledge of interaction recognition. Here, we introduce MARRI, a mechanism-aware multimodal framework that couples pretrained representations with domain knowledge by embedding learned opening penalties and hybridization gains into bidirectional cross-RNA attention. MARRI improves ROC–AUC and PR–AUC over the strongest baseline by 12.20% and 10.25%, respectively, and maintains strong discrimination for unseen RNA partnerships and interaction regions. Perturbation and repeat-context analyses identify prediction-relevant sequence regions. Integrating MARRI-derived interaction propensities with disease-associated expression profiles links molecular prediction to systems-level models of disease progression and network remodeling. MARRI thus illustrates a knowledge-guided strategy for coupling molecular foundation models with domain knowledge to support grounded prediction, interpretation and systems-level hypothesis generation.

## 1 Introduction

RNA–RNA interactions (RRIs) constitute a fundamental layer of post-transcriptional gene regulation, mediating molecular communication among coding and non-coding RNA species ^1,2^. Through direct base-pairing interactions, RNAs participate in diverse biological processes, including transcript processing, RNA stability, subcellular localization, and translational regulation. For example, microRNAs regulate gene expression by recognizing complementary sequences in target messenger RNAs, whereas long non-coding RNAs can interact with other RNA molecules to modulate transcript stability, processing, and regulatory complex formation ^1–3^. Advances in high-throughput experimental technologies, including CLASH, PARIS, SPLASH, and LIGR-seq, have substantially expanded our understanding of RNA interaction landscapes by enabling the systematic identification of intermolecular RNA contacts ^4–7^. Nevertheless, experimental detection remains limited by RNA abundance, cellular context, and assay coverage, leaving much of the RNA interactome unexplored. To help address this gap, we focus on RNA-pair interaction prediction, estimating whether two RNA sequence windows interact using their sequences, candidate local regions and biological annotations.

Computational prediction of RRIs has traditionally relied on thermodynamic models that evaluate RNA sequence complementarity and hybridization energetics. Methods such as RNAhybrid ^8^, RNAcofold ^9^, RNAplex ^10^, IntaRNA2^11^, RNAup ^12^, and RIsearch2^13^ identify potential interaction regions and estimate duplex stability, with several also accounting for the energetic costs of disrupting intramolecular RNA structures. These methods provide a physically grounded framework for predicting RNA interactions, but thermodynamic favorability alone cannot fully explain experimentally observed interactions. More recently, learning-based approaches, such as RIME ^14^, have sought to capture interaction-associated patterns directly from RNA sequences. However, sequence-based models such as RIME do not explicitly incorporate intramolecular RNA structural information into interaction scoring. Complementary regions may already participate in intramolecular base pairing, limiting their accessibility for intermolecular interactions ^15^. Predicting RRIs therefore requires considering both pairing potential and the structural availability of candidate interaction regions. The regulatory roles of interacting RNAs ^7^ and the positions of candidate sites within transcripts ^16^ provide additional biological context. Existing methods have not fully integrated these factors into RRI prediction.

Recent advances in RNA foundation models have enabled the learning of transferable representations for diverse biological applications. Models such as RNA-FM ^17^, RNAErnie ^18^, RiNALMo ^19^, RNA-MSM ^20^, and ERNIE-RNA^21^ have demonstrated the potential of large-scale pretraining to capture sequence, evolutionary, and structural features relevant to RNA function. Complementary advances in RNA secondary structure prediction, including RibonanzaNet ^22^, RNADiffFold ^23^, and trRNA2-SS ^24^, have further advanced data-driven modeling of RNA base-pairing patterns and secondary structures. Beyond structure prediction, pretrained RNA representations have been applied to a range of biological tasks. EvoRMD ^25^, SMRTnet ^26^ and ProRB ^27^ extend pretrained RNA representations to RNA modification prediction, RNA–small molecule binding and RNA–protein interaction tasks, respectively. Pretrained RNA foundation models provide rich representations of individual RNAs, but translating these representations into interaction predictions requires modeling the mechanistic constraints that govern intermolecular recognition. This raises a broader computational challenge: how to couple pretrained molecular representations with domain-specific mechanistic knowledge to support biologically grounded interaction prediction and interpretation.

MARRI addresses this challenge in the RNA setting by combining global and local RNA-FM representations with biological annotations, while embedding structural accessibility and intermolecular pairing directly into bidirectional axial cross-attention through learned opening penalties and hybridization gains. We evaluate MARRI through comparative benchmarks, component ablations and stringent held-out settings involving unseen RNA partners, RNA pairs and interaction loci. Beyond pair-level prediction, perturbation-based hotspot analysis and transposable-element enrichment characterize sequence regions and repeat contexts associated with predicted interactions. We further integrate fixed MARRI interaction propensities with disease-associated expression profiles in diabetic nephropathy and atherosclerosis, linking molecular interaction prediction to systems-level models of disease progression and network remodeling. Together, MARRI illustrates how domain knowledge can guide molecular foundation models across biological scales, from individual-molecule representation, through mechanism-aware interaction prediction and interpretation, to systems-level modeling of disease-associated network remodeling.

## 2 Results

We evaluated whether MARRI could combine accurate RNA-pair prediction with interpretable sequence-level evidence and informative downstream applications. First, we used comparative benchmarks, component ablations and held-out evaluations to assess predictive performance, the contributions of model components and generalization to unseen RNAs and interaction regions. Next, we examined the determinants underlying these predictions through hotspot perturbation, experimental structural comparisons and transposable-element enrichment analysis. Finally, we applied MARRI to diabetic nephropathy and atheroscle-rotic plaques to link MARRI predictions with disease-associated expression profiles to identify changes in candidate-network organization and prioritize RNA pairs for further investigation.

### 2.1 Overview of the framework and downstream analysis workflow

We developed MARRI, a mechanism-aware multimodal framework designed to predict RNA–RNA interactions (RRIs) by integrating sequence representations, intramolecular structural information and biological context (Fig. 1a). MARRI combines global and local RNA-FM representations to capture sequence context at complementary scales, using 256-nt windows alongside local candidate interaction regions. RibonanzaNet-SS base-pair confidence matrices are summarized into nucleotide-level structural features, while metadata embeddings encode RNA class and transcript-region annotations. To integrate these complementary inputs, MARRI employs bidirectional axial cross-attention that incorporates learned opening penalties and pairwise hybridization gains. The opening penalties represent structural constraints on potential interaction sites, whereas hybridization gains provide learned pairing-related signals between nucleotides from the two RNA partners. Together, these components enable the model to account for local structural accessibility and intermolecular pairing when predicting RNA interactions. MARRI outputs an RNA-pair interaction probability and a nucleotide-pair evidence map, supporting both interaction prediction and sequence-level interpretation. A detailed architecture is provided in Supplementary Fig. S1.

**Figure 1:**
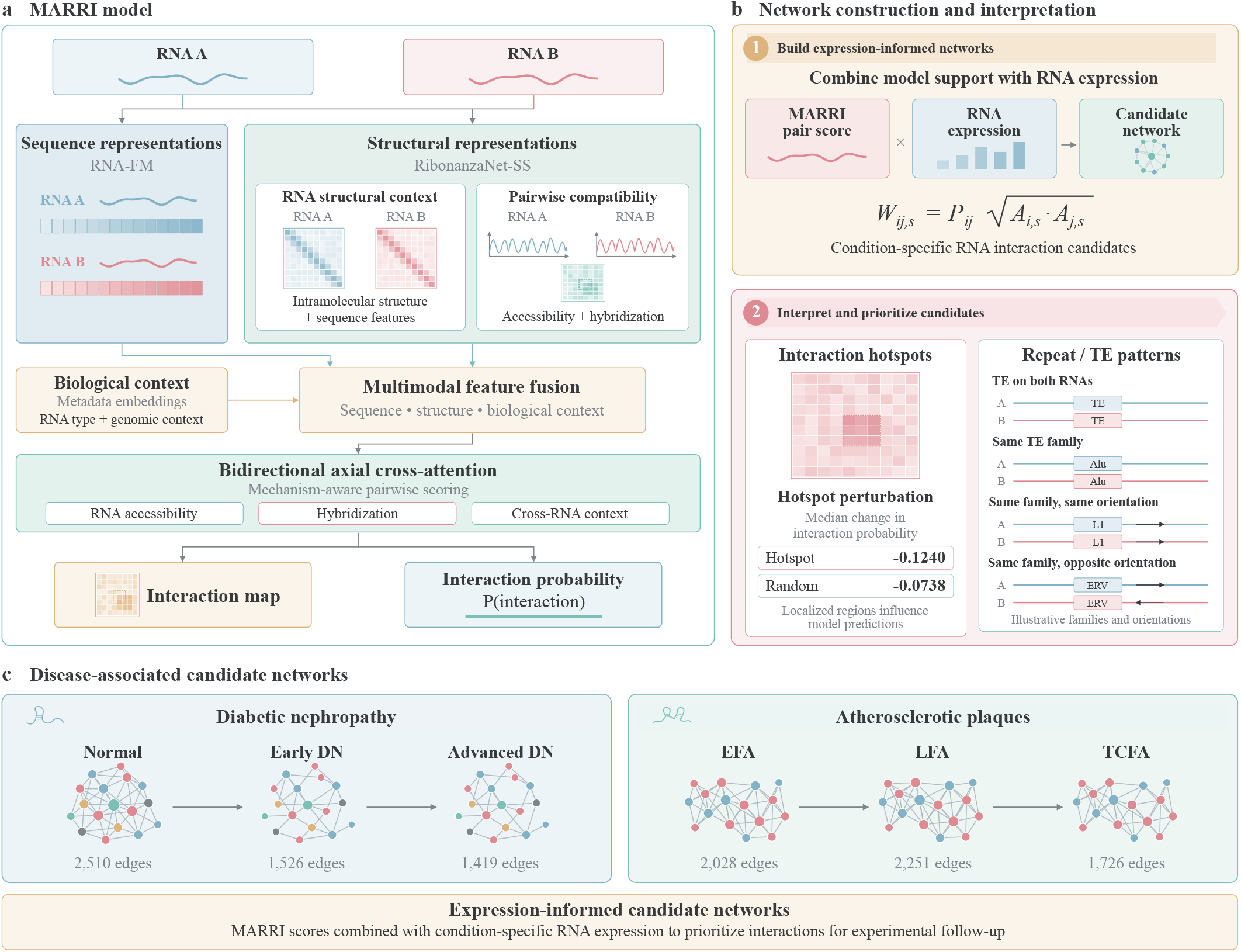
MARRI overview, evaluation and biological applications. **a**, RNA-FM sequence representations, RibonanzaNet-SS structural features and RNA metadata are integrated through bidirectional axial cross-attention. The model produces an RNA-pair interaction probability and a nucleotide-pair evidence map. **b**, downstream analysis combines fixed MARRI scores with RNA expression to construct candidate networks. Hotspot perturbation and TE enrichment examine local score dependence and repeat-associated sequence context. **c**, candidate networks in diabetic nephropathy and atherosclerotic plaques. Edge counts indicate backbone sizes at a shared threshold within each disease analysis.

Beyond pair-level prediction, MARRI outputs supported hotspot analysis and the identification of repeat-associated sequence patterns (Fig. 1b). Integration with disease-associated expression profiles enabled construction of candidate RNA interaction networks in diabetic nephropathy and atherosclerotic plaques (Fig. 1c).

### 2.2 MARRI performance relative to existing methods

To evaluate the predictive performance of MARRI, we benchmarked it against representative learning-based and thermodynamics-based methods using a common held-out test set (Table 1). MARRI consistently outperformed all evaluated comparators in both ROC–AUC and PR–AUC, achieving (74.27 *±* 0.95)% and (72.24 *±* 1.15)%, respectively, across three independently trained models. Compared with RIME ^14^, the learning-based baseline, MARRI improved ROC–AUC by 12.20 percentage points and PR–AUC by 10.25 percentage points. These improvements indicate that MARRI more effectively distinguishes labeled interacting from non-interacting RNA pairs within the evaluated benchmark.

**Table 1.** Comparison of MARRI with learning-based and physics-based predictors. Values are percentages. MARRI and RIME results are means *±* sample standard deviations across seeds 42, 43 and 44. Physics-based methods were evaluated once.

| Method | Type | ROC–AUC | PR–AUC | Accuracy | Precision | Recall | F1-score |
|---|---|---|---|---|---|---|---|
| MARRI (ours) | Learning-based | **74.27 ± 0.95** | **72.24 ± 1.15** | **67.42 ± 0.83** | **65.38 ± 1.40** | 73.47 ± 1.36 | **69.17 ± 0.26** |
| RIME | Learning-based | 62.07 ± 0.38 | 61.99 ± 0.42 | 58.05 ± 0.47 | 57.26± 0.70 | 61.77 ± 0.49 | 59.43 ± 0.58 |
| IntaRNA2 | Physics-based | 54.14 | 53.99 | 49.73 | 49.71 | 99.91 | 66.39 |
| RNAplex | Physics-based | 55.97 | 57.82 | 49.95 | 49.96 | 99.92 | 66.61 |
| RNAcofold | Physics-based | 53.56 | 55.97 | 49.94 | 49.95 | 99.91 | 66.60 |
| RNAhybrid | Physics-based | 56.07 | 57.96 | 49.95 | 49.96 | 99.92 | 66.61 |
| RNAup | Physics-based | 52.55 | 53.43 | 50.06 | 50.01 | 99.88 | 66.65 |
| RIsearch2 | Physics-based | 58.46 | 61.83 | 50.02 | 49.99 | **99.97** | 66.65 |

The performance differences were particularly evident when compared with thermodynamics-based approaches, including IntaRNA2, RNAplex, RNAcofold, RNAhybrid, RNAup and RIsearch2^8–13^. Although these methods achieved near-complete recall at their evaluated operating points, their precision remained close to 50%, indicating limited discrimination between positive and negative RNA pairs under the benchmark conditions. Even RIsearch2, the strongest thermodynamics-based comparator in terms of ROC–AUC and PR–AUC, achieved only 58.46% and 61.83%, respectively. In contrast, MARRI achieved a more balanced precision–recall profile, with precision of 65.38%, recall of 73.47% and an F1-score of 69.17%. Together, these results demonstrate improved predictive discrimination by MARRI over both sequence-learning and thermodynamics-based approaches in the evaluated test set.

### 2.3 Contributions of model components to predictive performance

To assess the contributions of individual components, we systematically ablated local interaction-window representations, biological metadata, RibonanzaNet-SS structural features and pairwise hybridization gain *H*_*ij*_, retraining each variant under the same protocol across three random seeds (Table 2).

**Table 2.**
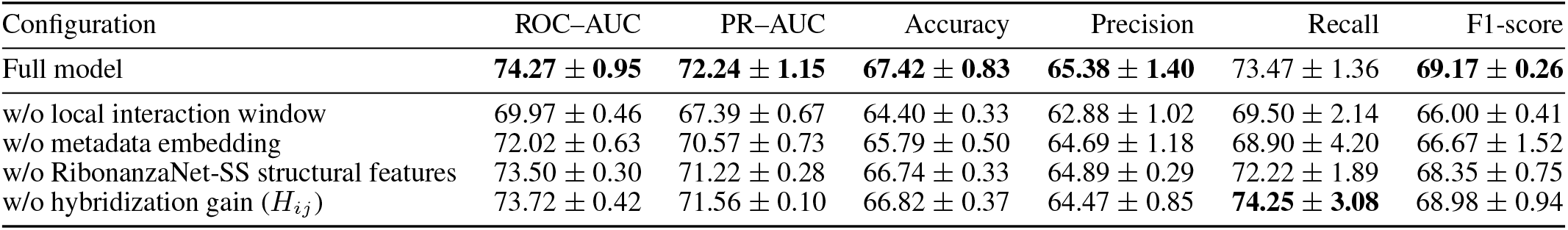
Performance after individual component ablations. The full model fine-tuned RNA-FM, froze RibonanzaNet-SS and used dynamic negative resampling. Values are percentages, reported as means *±* sample standard deviations across seeds 42, 43 and 44.

| Configuration | ROC–AUC | PR–AUC | Accuracy | Precision | Recall | F1-score |
|---|---|---|---|---|---|---|
| Full model | **74.27 ± 0.95** | **72.24 ± 1.15** | **67.42 ± 0.83** | **65.38 ± 1.40** | 73.47 ± 1.36 | **69.17 ± 0.26** |
| w/o local interaction window | 69.97 ± 0.46 | 67.39 ± 0.67 | 64.40 ± 0.33 | 62.88 ± 1.02 | 69.50 ± 2.14 | 66.00 ± 0.41 |
| w/o metadata embedding | 72.02 ± 0.63 | 70.57 ± 0.73 | 65.79 ± 0.50 | 64.69 ± 1.18 | 68.90 ± 4.20 | 66.67 ± 1.52 |
| w/o RibonanzaNet-SS structural features | 73.50 ± 0.30 | 71.22 ± 0.28 | 66.74 ± 0.33 | 64.89 ± 0.29 | 72.22 ± 1.89 | 68.35 ± 0.75 |
| w/o hybridization gain (*H*<sub>*ij*</sub>) | 73.72 ± 0.42 | 71.56 ± 0.10 | 66.82 ± 0.37 | 64.47 ± 0.85 | **74.25 ± 3.08** | 68.98 ± 0.94 |

Explicitly incorporating local interaction-window representations improved ROC–AUC and PR–AUC by 4.30 and 4.85 percentage points relative to the variant without this branch. This gain highlights the importance of separately encoding candidate interaction regions alongside broader sequence context. MARRI’s explicit global–local fusion extends the sequence-embedding and contact-matrix design used by RIME ^14^. Incorporating biological metadata improved ROC–AUC and PR–AUC by 2.25 and 1.67 percentage points, respectively. These improvements demonstrate the predictive value of RNA-class and transcript-region annotations, which are not explicitly supplied to the sequence-based RIME model.

Incorporating RibonanzaNet-SS-derived structural features improved ROC–AUC and PR–AUC by 0.77 and 1.02 percentage points relative to the corresponding ablated variant. This result supports explicitly providing intramolecular structural information beyond learned sequence representations. Such information is not directly included in RIME’s inputs and enables MARRI to incorporate structural constraints into accessibility-aware pairwise scoring. Including the hybridization gain *H*_*ij*_ yielded additional improvements of 0.55 and 0.68 percentage points over its ablated variant, supporting context-dependent modeling of intermolecular pairing potential.

Together, these results support extending sequence-based RRI prediction through explicit global–local encoding, biological annotations and structure-informed interaction scoring. Dynamic negative resampling also yielded small but consistently positive improvements over fixed negative sampling across four backbone configurations (Supplementary Table S2).

### 2.4 Generalization across unseen RNAs and interaction regions

To assess MARRI’s generalizability beyond previously observed interactions, we evaluated three non-random held-out strategies designed to prevent information sharing at the gene, RNA-pair and interaction-locus levels (Supplementary Table S5). All three strategies started from the same pool of positive interactions and targeted an 80:10:10 training, validation and test split, with each gene group, unordered RNA pair or interaction-locus group assigned to a single partition in the corresponding evaluation. These settings progressively examine distinct forms of generalization, from recognizing new interaction regions and RNA partnerships to predicting interactions involving entirely unseen genes. MARRI maintained predictive discrimination under all three settings, achieving ROC–AUC values of 75.02% for locus-held-out, 74.22% for pair-held-out and 66.30% for gene-held-out evaluation.

Performance varied across the three generalization settings, with the strongest discrimination observed for unseen interaction loci and RNA partnerships. The corresponding PR–AUC values were 33.51% and 32.23%, compared with 17.50% for unseen genes. Gene-held-out evaluation also yielded the lowest F1-score (23.01%), compared with 35.50% and 35.81% for pair-and locus-held-out evaluation, respectively. These findings indicate that MARRI can distinguish candidate interactions involving previously unseen RNA partnerships and local regions, while generalization to entirely unseen genes remains more challenging.

### 2.5 Model interpretability and sequence context

To investigate the sequence-level basis of MARRI predictions, we combined hotspot perturbation analysis with transposable-element (TE) enrichment analysis. These complementary approaches assessed whether model-identified interaction regions influence prediction scores and whether predicted interactions exhibit biologically relevant sequence patterns.

#### 2.5.1 Local hotspot perturbation supports MARRI score faithfulness

To evaluate whether model-identified hotspots contribute to interaction predictions, we compared the effects of hotspot disruption with composition-matched substitutions at other positions within the same RNA pairs. Across 182 high-confidence positive test-set pairs, hotspot disruption produced significantly larger decreases in predicted interaction probability than matched random-site substitutions (median Δ*P* = −0.124 versus −0.074; one-sided paired Wilcoxon test, *P* = 1.89 ×10^−9^; Fig. 2a–d).

**Figure 2:**
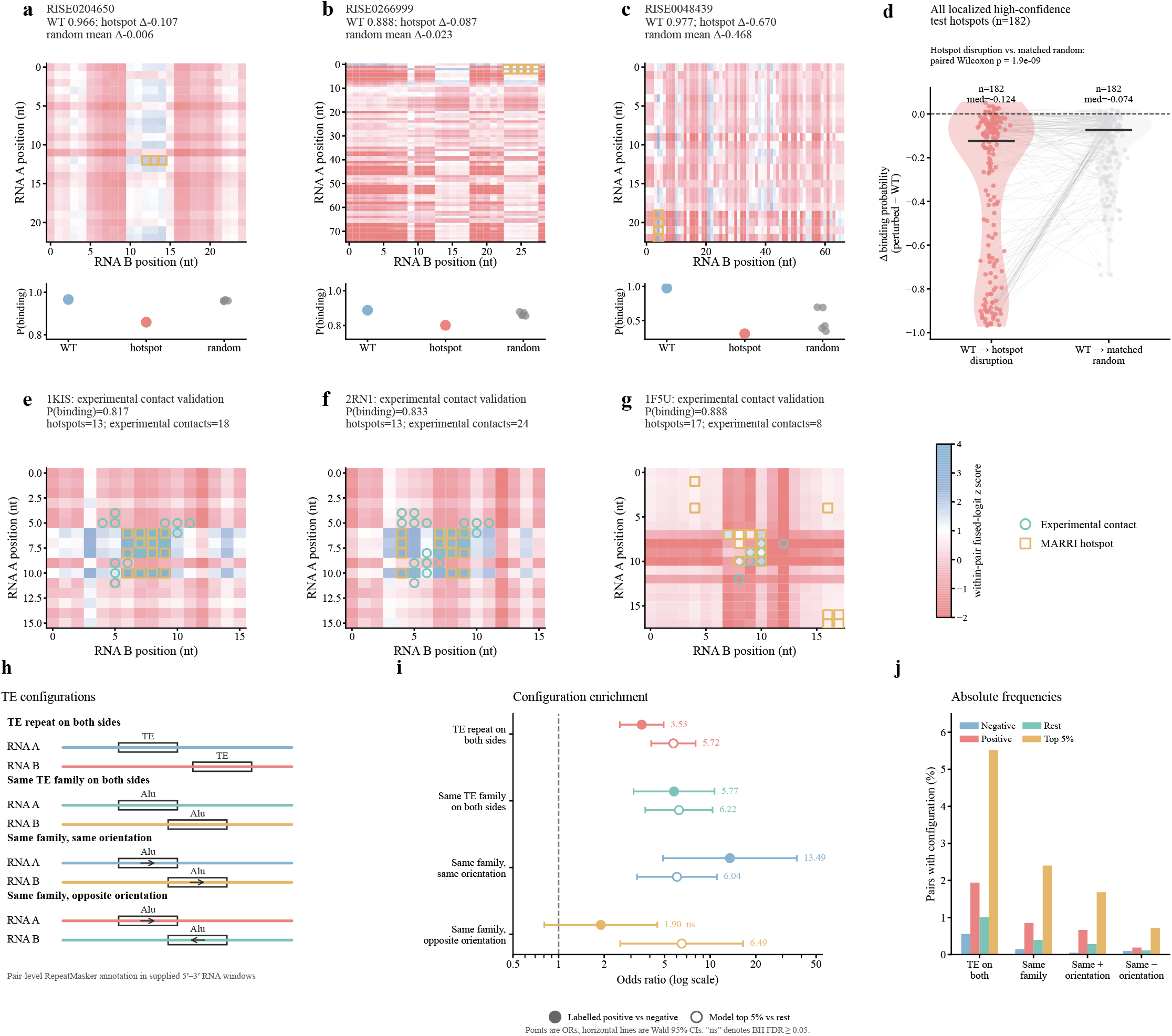
MARRI hotspot analysis and TE enrichment. **a–c**, representative positive test-set pairs. Heatmaps show within-pair standardized fused logits; yellow squares mark hotspots. Lower plots show probabilities for wild-type (WT), hotspot-disruption and matched random-site inputs. **d**, paired probability changes across 182 candidates. Each line connects the hotspot effect with the mean of two to five matched random-control effects. Horizontal bars indicate medians. **e–g**, hotspots and experimental contact cells for 1KIS, 2RN1 and 1F5U ^28–30^. Cyan circles mark experimental contacts and yellow squares mark hotspots. Exact contact-cell F1-scores were 0.323, 0.324 and 0.400, respectively. **h**, RepeatMasker configurations in the supplied 5^′^ –3^′^local RNA windows. **i**, odds ratios for positive versus negative pairs and top-5%-score versus remaining pairs. Horizontal lines indicate Wald 95% confidence intervals; “ns” denotes Benjamini–Hochberg FDR 0≥.05. **j**, the corresponding absolute configuration frequencies.

These results indicate that the identified hotspots capture sequence regions that exert a greater influence on MARRI predictions than matched control regions.

To examine their correspondence with experimentally observed RNA interactions, we further compared MARRI hotspots with contact maps from three RNA complexes: 1KIS, 2RN1 and 1F5U ^28–30^. The resulting exact contact-cell F1-scores were 0.323,0.324 and 0.400, respectively (Fig. 2e–g), indicating partial correspondence between model-identified hotspots and experimental interaction interfaces. Although these comparisons do not establish that individual hotspots represent physical contact sites, they provide an initial structural consistency assessment of MARRI’s sequence-level interpretations.

#### 2.5.2 TE repeat enrichment

We next examined whether transposable-element (TE) configurations were associated with experimentally labeled interactions and high-scoring MARRI predictions. Using RepeatMasker ^31^, we annotated local RNA windows and assessed the occurrence of TE sequences in both interacting partners, shared TE families and shared-family repeats with matching or opposing transcript orientations (Fig. 2h). Enrichment was evaluated by comparing 7,982 positive and 8,076 negative test-set pairs, as well as 833 top-scoring pairs with the remaining test-set pairs, using Fisher’s exact tests with Benjamini–Hochberg correction.

TE annotations in both RNA partners were more frequent among positive than negative pairs (1.94% versus 0.56%; OR = 3.53, FDR = 2.92 × 10^−15^) and were further enriched among the top 5% of MARRI predictions (5.52% versus 1.01%; OR = 5.72, FDR = 8.52 × 10^−17^; Fig. 2i,j). Shared TE families showed similar enrichment in both comparisons (OR = 5.77 and 6.22, respectively), suggesting that repeat-associated sequence relationships are represented among both labeled interactions and model-prioritized candidates.

Repeat orientation further distinguished these associations. Shared-family repeats with matching orientations showed the strongest enrichment among positive versus negative pairs (OR = 13.49, FDR = 6.19 × 10^−12^) and were also enriched among high-scoring predictions (OR = 6.04, FDR = 1.27 × 10^−6^). Opposite-orientation repeats were enriched among high-scoring pairs (OR = 6.49, FDR = 0.00103), although this association involved only six pairs and was not significant in the positive-versus-negative comparison (FDR = 0.149).

Together, these findings identify repeat-associated sequence configurations enriched among labeled RNA interactions and MARRI-prioritized candidates. Previous studies have implicated complementary Alu-derived sequences in STAU1-mediated mRNA decay ^32^ and enhancer–promoter communication ^33^, providing biological precedents for repeat-mediated RNA recognition. Although the observed enrichments do not establish direct TE-mediated pairing in individual predictions, they highlight repeat-containing regions as candidates for future experimental investigation.

### 2.6 Disease-associated RNA–RNA network remodeling

We next asked whether MARRI could extend beyond static RNA–RNA interaction prediction to identify candidate interaction networks that vary across disease contexts. We therefore combined fixed MARRI interaction scores with RNA expression availability and examined network organization in two distinct disease settings: diabetic nephropathy (DN) and atherosclerosis. Rather than interpreting these expression-conditioned networks as direct measurements of physical interactions, we used them to identify interaction candidates whose network prominence changes across disease states.

#### 2.6.1 Diabetic nephropathy

We first examined Normal, Early DN and Advanced DN expression backgrounds from GSE142025^34^. Among database-supported RNA pairs recorded in RNAInter or NPInter ^35,36^, MARRI prioritized 24,452 of 62,613 scorable pairs (39.1%) at *P*_*ij*_ ≥ 0.8 (Fig. 3b). The reference scaffold further restricted candidates to pairs scored in all three groups and retained only the high-score backbone (Fig. 3a).

**Figure 3:**
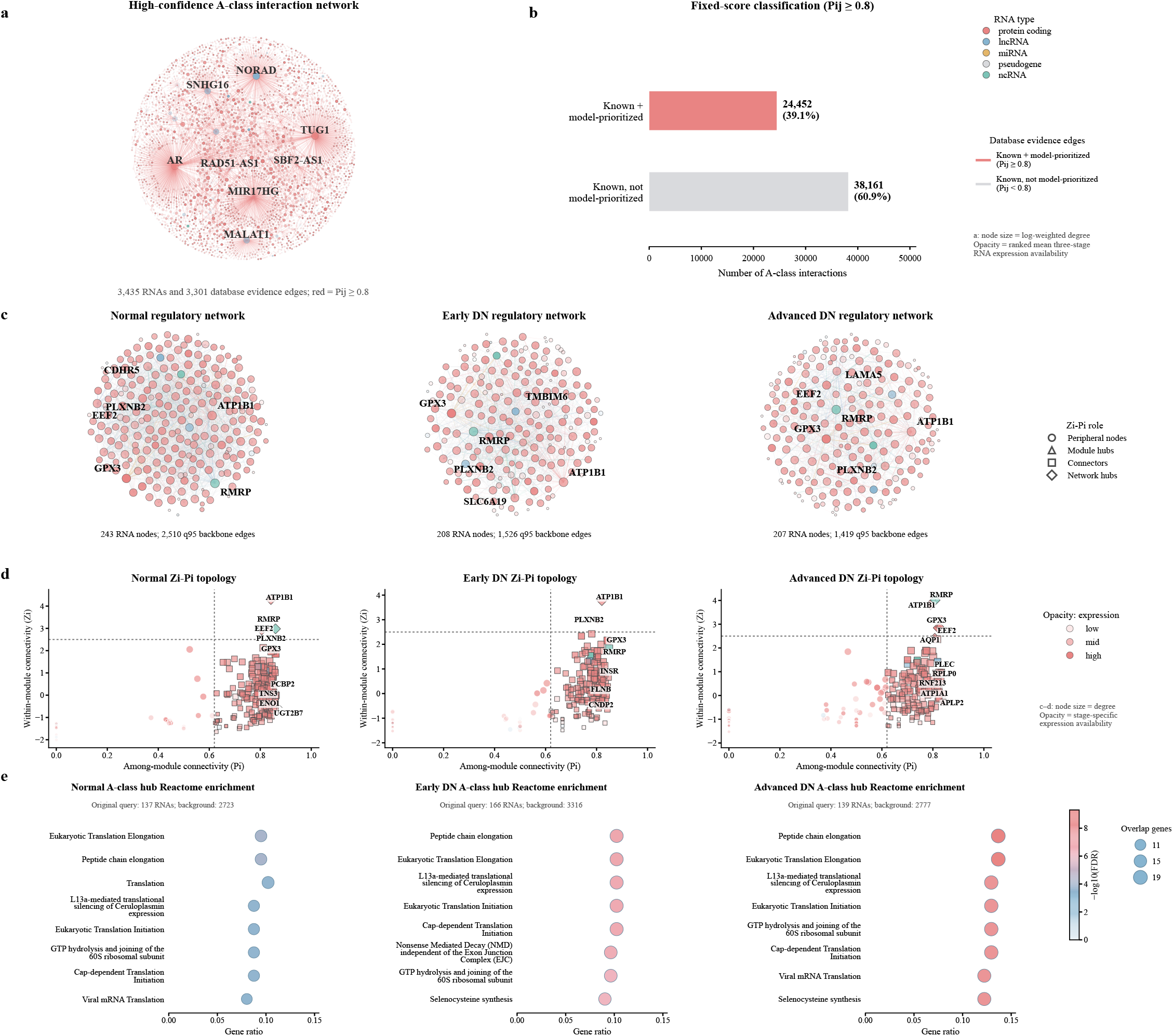
Evidence-supported candidates, network organization and functional enrichment in diabetic nephropathy. **a**, high-confidence A-class scaffold with 3,435 RNA nodes and 3,301 database-supported edges. Node color denotes RNA type, size denotes log-transformed weighted degree and opacity denotes ranked mean expression availability across the three groups. Red edges have *P*_*ij ≥*_0.8; labels identify highly connected candidates. **b**, scores for all 62,613 scorable database-supported pairs. At *P*_*ij ≥*_0.8, 24,452 pairs (39.1%) were prioritized and 38,161 (60.9%) fell below the threshold. **c**, exploratory B-class backbones for Normal, Early DN and Advanced DN, extracted at a shared pooled 95th-percentile edge-weight threshold. Node color denotes RNA type, size denotes degree and opacity denotes stage-specific expression availability. **d**, weighted Zi–Pi roles for the corresponding B-class backbones. *Z*_*i*_ measures within-module connectivity and *P*_*i*_ measures participation across modules; dotted lines indicate *Z*_*i*_ = 2.5 and *P*_*i*_ = 0.62. **e**, Reactome enrichment of the top-5% weighted-degree A-class RNAs, using each stage’s network as background. The eight leading significant pathways are displayed. Horizontal position denotes pathway-overlap genes divided by input RNAs; bubble area indicates overlap count and color indicates *−*log_10_(FDR).

The exploratory B-class networks contained fewer high-weight candidate edges in the DN groups. At a shared pooled 95th-percentile edge-weight threshold, the Normal backbone contained 243 RNAs and 2,510 edges. Early DN contained 208 RNAs and 1,526 edges, whereas Advanced DN contained 207 RNAs and 1,419 edges (Fig. 3c). Changes in connectivity were accompanied by shifts in network roles. Zi–Pi analysis identified 212 cross-module connectors in Normal, decreasing to 163 in Early DN and 141 in Advanced DN; the corresponding proportions declined from 87.2% to 78.4% and 68.1%, respectively (Fig. 3d). Together, these results indicate progressively fewer high-weight candidate connections and fewer RNAs linking network modules across DN expression backgrounds.

Network remodeling also highlighted individual RNAs with independent links to renal disease biology. ATP1B1 remained a prominent network hub across disease stages, consistent with human single-cell multi-omic evidence of disease-associated chromatin accessibility changes near ATP1B1 in diabetic kidney disease ^37^. RMRP changed in network prominence and is particularly notable because it has been experimentally implicated in DN through an RMRP–miR-1a-3p–JunD regulatory axis that promotes mesangial-cell proliferation and fibrosis ^38^. GPX3 likewise showed altered network prominence; as an extracellular antioxidant enzyme synthesized largely by the kidney, GPX3 has been linked experimentally to oxidative and thrombotic complications of chronic kidney disease ^39^. These examples illustrate how disease-conditioned MARRI networks can prioritize candidate RNAs whose altered network roles converge with independent disease evidence.

Despite these topological changes, highly connected RNAs showed a recurrent functional signature across disease stages. Reactome enrichment of the top-5% weighted-degree A-class RNA sets identified translation-related processes in Normal, Early DN and Advanced DN, including translation initiation and elongation, peptide-chain elongation, cap-dependent initiation, 60S-subunit joining and L13a-mediated translational silencing (Fig. 3e). Additional enriched terms included nonsense-mediated decay and selenocysteine synthesis (linked processes given that the codon for selenocysteine is a stop codon ^40,41^), linking the network signal to transcript surveillance and specialized protein synthesis. These recurrent annotations are consistent with reported alterations in protein-translation programs in human and experimental diabetic kidney disease ^42^. Thus, DN progression was associated with substantial remodeling of candidate RNA-network topology while retaining a shared translation-related functional signature among highly connected RNAs.

#### 2.6.2 Atherosclerosis

We next examined whether disease-associated network remodeling could be observed in an independent biological setting. We analyzed carotid plaques classified as early fibroatheroma (EFA; *n* = 4), late fibroatheroma (LFA; *n* = 4) or thin-cap fibroatheroma (TCFA; *n* = 5) ^43^. TCFA plaques showed a distinct transcriptional state relative to LFA, with 3,235 differentially expressed genes at FDR < 0.05 and | log_2_ FC| > 1. This state was characterized by increased macrophage-and antigen-presentation-associated expression, reduced smooth-muscle-associated expression and enrichment of immune and innate immune pathways (Fig. 4a,b,d), consistent with established immune-cell and smooth-muscle phenotypic changes in human atherosclerotic plaques ^44,45^.

**Figure 4:**
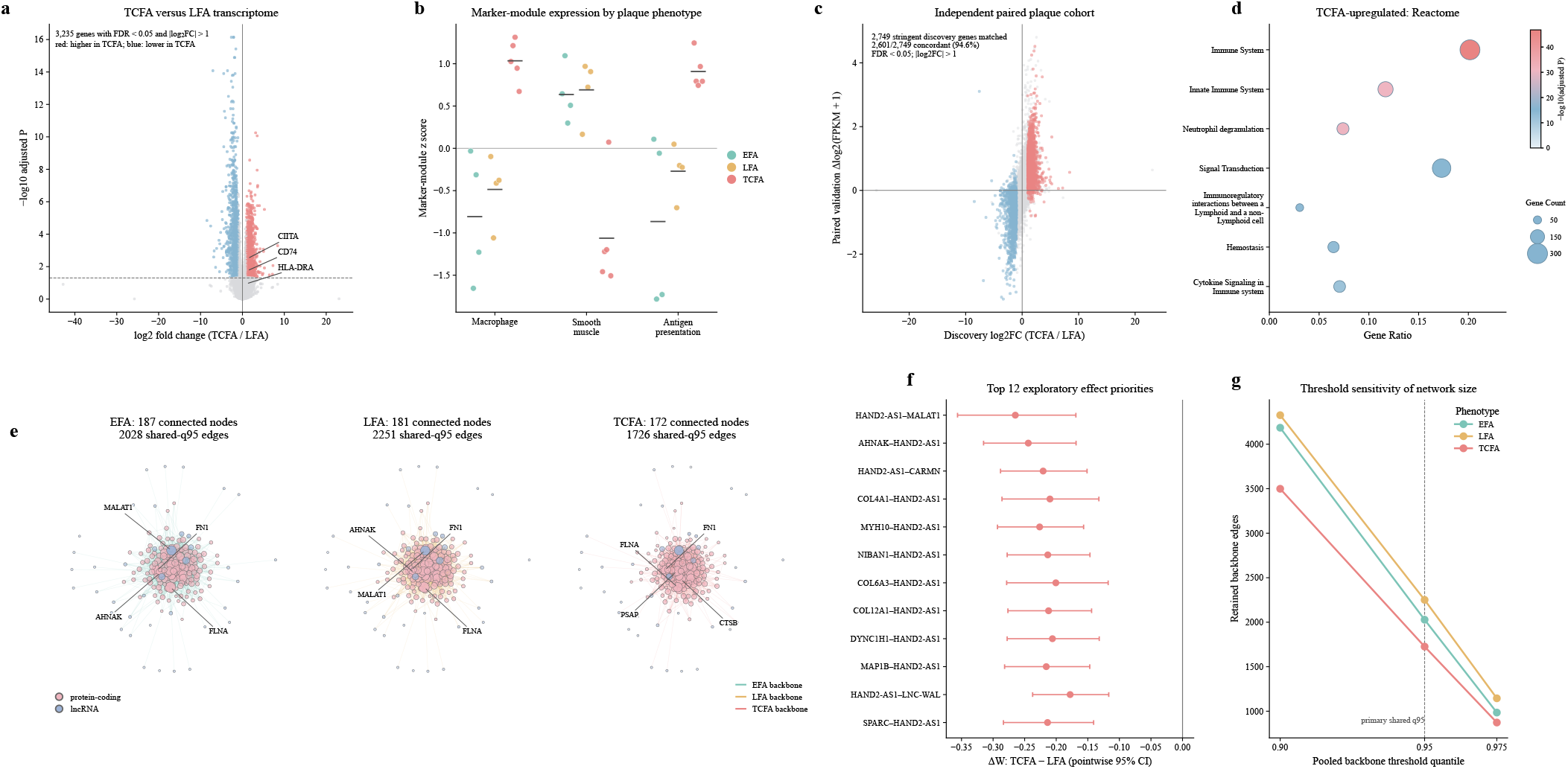
Expression context and candidate-network remodeling across plaque phenotypes. **Expression context (a–d)** describes transcriptional differences and their functional associations. **a**, TCFA–LFA differential expression. Red and blue points meet FDR *<* 0.05 and |log_2_ FC |*>* 1, with higher expression in TCFA and LFA, respectively. **b**, prespecified marker-module scores; points indicate samples and horizontal bars indicate means. **c**, expression-effect concordance with paired unstable–stable regions in GSE120521^46^. Colored points mark the 2,749 matched stringent discovery genes. **d**, Reactome enrichment of stringent TCFA-upregulated genes. Position denotes gene ratio, bubble area denotes gene count and color denotes −log_10_ adjusted *P*. **MARRI network application (e–g)** examines candidate connectivity, individual pairs and threshold sensitivity. **e**, EFA, LFA and TCFA backbones at a shared pooled 95th-percentile threshold. Node color denotes RNA class, size denotes degree and opacity denotes phenotype-level mean availability. **f**, twelve effect-prioritized candidate edges. Points show TCFA–LFA weight differences and whiskers show pointwise 95% bootstrap intervals. **g**, retained edge counts at prespecified pooled quantiles.

The transcriptional pattern was independently supported using paired stable and unstable plaque regions from GSE120521^46^.Among 2,749 matched genes meeting the stringent discovery criteria, 2,601 (94.6%) changed in the same direction, while effect sizes across all 21,000 matched genes showed a Spearman correlation of *ρ* = 0.679 (Fig. 4c). This cross-dataset concordance provided an independent expression context in which to interpret the MARRI-derived network changes. Within a common panel of 300 RNAs, expression-conditioned MARRI networks showed reduced candidate connectivity in TCFA. At the shared pooled 95th-percentile threshold, EFA, LFA and TCFA retained 2,028, 2,251 and 1,726 edges, respectively, corresponding to a 23.3% reduction in TCFA relative to LFA (Fig. 4e). The ordering LFA > EFA > TCFA persisted across pooled thresholds from the 90th to the 97.5th percentile (Fig. 4g), indicating that the observed reduction was not specific to a single network threshold.

At the individual-interaction level, the TCFA–LFA comparison prioritized decreased candidate-edge weights centered on *HAND2-AS1*, which suppresses lipid accumulation and foam cell formation in atheroma ^47^(Fig. 4f). The *HAND2-AS1*–*MALAT1* pair had a high fixed MARRI interaction score of 0.942, whereas its mean expression-conditioned weight decreased from 0.507 in LFA to 0.243 in TCFA (Δ*W* = −0.265; FDR = 0.0535). The *HAND2-AS1*–*CARMN* interaction showed a similar decrease (Δ*W* = −0.221; FDR = 0.0493). Both MALAT1 and CARMN have been associated with atherogenesis, although the mechanisms are complex and incomplete at this time ^48,49^. Thus, the global reduction in candidate-network connectivity could be resolved to specific RNA pairs that may warrant targeted mechanistic investigation.

Across DN and atherosclerosis, expression-conditioned MARRI networks, therefore, revealed disease-associated changes at complementary scales. Both settings showed reduced high-weight candidate connectivity in more advanced disease states, while network topology further prioritized disease-relevant RNAs and individual interaction candidates. These analyses illustrate how MARRI can translate sequence-derived interaction predictions into context-dependent hypotheses about RNA-network organization and disease-associated remodeling.

## 3 Discussion

RNA–RNA interactions (RRIs) are governed by an interplay among factors, including sequence complementarity, RNA structural accessibility, and biological context. MARRI addresses this complexity through a mechanism-aware multimodal framework that integrates pretrained RNA representations, intramolecular structural features, and biological annotations. By incorporating learned opening penalties and pairwise hybridization gains into bidirectional cross-RNA attention, MARRI explicitly models both structural constraints and intermolecular pairing. Its improved predictive performance over the evaluated learning-based and thermodynamics-based methods underscores the value of integrating complementary sources of information for RRI prediction. MARRI extends beyond binary pair-level classification by connecting predictive evidence with local sequence interpretability and disease-associated network analysis.

Our analyses offer insights into the important determinants of accurate RRI prediction. Local interaction-window representations contributed the largest performance gains, underscoring the importance of resolving candidate interaction regions within broader RNA sequence contexts. RNA-class and transcript-region annotations provided additional predictive value, highlighting the relevance of biological context and its complementarity with sequence representations. Structural features and pairwise hybridization gains yielded yet complementary improvements, justifying their integration within the overall framework. Generalization experiments further showed that MARRI maintained strong predictive discrimination for unseen RNA partnerships and interaction regions, while interactions involving entirely unseen genes remain more challenging. Together, these findings indicate that while that local sequence patterns and RNA-specific biological contexts are important for interaction recognition, learning representations that transfer across distinct RNA identities remains an open challenge.

Beyond predictive performance, MARRI provides opportunities to investigate the sequence features governing RNA recognition. Local hotspot perturbation revealed regions whose disruption had a greater effect on predicted interaction probabilities than composition-matched control substitutions, confirming the importance of model-identified hotspots. Their partial overlap with experimentally resolved contacts across three RNA complexes further indicates the correspondence between model-highlighted regions and physical interaction interfaces. Complementary transposable-element (TE) analyses identified repeat-associated configurations enriched among experimentally RNA interactions and high-scoring predictions. These findings are consistent with the roles of Alu-derived sequences in STAU1-mediated mRNA decay and enhancer–promoter communication, as well as the broader contribution of TEs to lncRNA sequence organization ^32,33,50^. Our hotspot and repeat-context analyses connect MARRI’s predictions with biologically relevant sequence features and generate testable hypotheses for investigating RNA recognition mechanisms.

Applying MARRI to diabetic nephropathy and atherosclerosis illustrates how sequence-informed interaction predictions can be integrated with disease-associated transcriptomic profiles to investigate candidate RNA interaction networks. In diabetic nephropathy, expression-conditioned networks exhibited fewer high-weight candidate interactions and cross-module connections, while translation-related functional enrichment was observed across disease states, pointing to a potential link between candidate RNA interaction patterns and translational regulation. In atherosclerotic plaques, shifts in candidate network connectivity coincided with immune-associated expression changes, where altered interaction weights involving *HAND2-AS1* prioritized RNA-centered candidates for further investigation. These analyses demonstrate the potential of combining predicted interaction propensity with disease-specific expression profiles to identify regulatory relationships that may not be apparent from transcript abundance alone. More broadly, this framework provides a systematic strategy for prioritizing disease-relevant RNA interactions and generating testable mechanistic hypotheses.

These findings highlight several promising avenues for future work. Expanding datasets to cover more diverse RNA partners could further improve generalization across biological contexts. Integrating cell-type-resolved expression profiles and orthogonal modalities may extend MARRI toward finer-grained interaction analysis. Additionally, post-transcriptional RNA modifications, such as m^6^A, may reshape RNA structure and interaction potential ^51,52^; epitranscriptomic modifications into future iterations of MARRI could help characterize how modification states reshape RNA–RNA interaction landscapes. Experimentally testing model-identified hotspots, repeat-associated sequences, and prioritized disease-related RNA pairs will also be essential to establish their functional roles. Ultimately, by connecting mechanism-aware interaction prediction with sequence-level interpretability and disease-associated network analysis, MARRI provides a framework for advancing our understanding of RNA recognition and guiding targeted experimental discovery.

## 4 Methods

### 4.1 Data preparation

#### 4.1.1 Data sources and sample construction

We curated experimentally supported human RNA–RNA interactions from RISE and RNAInter ^35,53^. Positive examples required valid genomic coordinates for both interacting segments. To reduce redundancy from fragmented assay records, we applied direction-independent deduplication based on fragment positions. Each fragment was assigned a chromosome, strand and center-coordinate signature using 50-nt fuzzy bins. We sorted the two signatures to obtain an undirected interaction identifier and collapsed records sharing that identifier into one positive interaction locus.

Using bedtools ^54^, we expanded each fragment into a 256-nt window centered on the annotated interaction region wherever possible. Fragment lengths had a 99th percentile of approximately 70 nt and a maximum below 81 nt. The expanded windows therefore retained the interaction region and flanking sequence for structural modeling. The two model inputs were

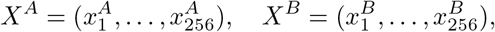

where *X*^*A*^ and *X*^*B*^ denote the context windows for RNA A and RNA B.

#### 4.1.2 Negative sample construction

We constructed a biotype-matched negative pool because validated non-interacting pairs were generally unavailable. Coordinate-boundary filtering and exact deduplication yielded 496,686 records, more than five times the positive-set size. Major RNA-biotype pairings were matched to the positive-set distribution to reduce class-based shortcuts.

The pool contained three negative categories. Partner-swapped negatives (PN; 50%) exchanged partners between positive interactions while preserving biotype constraints. They combined regions with individual interaction evidence into alternative pairings. Local-region negatives (NP; 25%) retained the RNA partners but shifted windows by 500–5000 nt along their transcripts. They contrasted annotated interaction regions with alternative regions of the same RNA pair. Background negatives (NN; 25%) randomly paired RNAs and sampled local windows to represent transcriptome-level alternatives.

We removed exact duplicates using the chromosome, start and end coordinates of both windows. At each training epoch, negatives were dynamically sampled to maintain approximately equal numbers of positive and negative pairs while retaining the PN:NP:NN proportions.

Before inference, sequences were uppercased and thymine was converted to uracil. A pair was retained only when both 256-nt context windows contained exclusively A, C, G and U, as required by RibonanzaNet-SS. This N-clean filter excluded ambiguous bases and other non-canonical characters and was applied during both training and downstream inference.

### 4.2 Architecture

MARRI integrates sequence context, RNA metadata, local candidate regions and intramolecular structural features. Its four components are multimodal encoding, internal mechanistic branches, accessibility-aware axial cross-attention and an RNA-pair classifier.

#### Multimodal sequence and metadata encoding

A shared RNA-FM encoder ^17^ generated contextual nucleotide representations for both RNA partners:

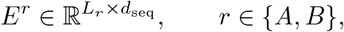

where *L*_*r*_ is the input-window length and *d*_seq_ = 640. We denote the feature vector at nucleotide position *i* by 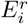. Global inputs were 256-nt context windows centered on candidate interaction regions. Local inputs retained the selected regions at their original lengths. The same encoder processed global and local inputs from both RNAs.

For each input RNA window *r* ∈ *{A, B}*, let *t*^*r*^ denote its RNA-type category and *c*^*r*^ its assigned transcript-region category. Two learnable embedding functions mapped these categorical annotations to vectors, which were concatenated into a window-level metadata representation:

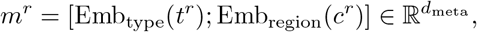

where *d*_meta_ is the combined embedding dimension and [*·*; *·*] denotes concatenation. At nucleotide position *i*, the sequence representation 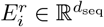 was concatenated with the same metadata vector:

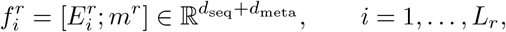

where *L*_*r*_ is the input-window length. Thus, sequence features varied across nucleotide positions, whereas the metadata vector was shared across all positions within the window. This encoding procedure was applied to both global and local inputs.

#### Intramolecular structural features

We used RibonanzaNet-SS, a secondary-structure-fine-tuned RibonanzaNet model initialized from publicly released weights ^22^. It remained frozen during RNA-FM fine-tuning and generated a continuous intramolecular pair-confidence matrix for each RNA.

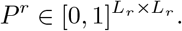

Before aggregation, we masked near-diagonal entries:

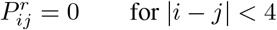

We then calculated four descriptors for each nucleotide:

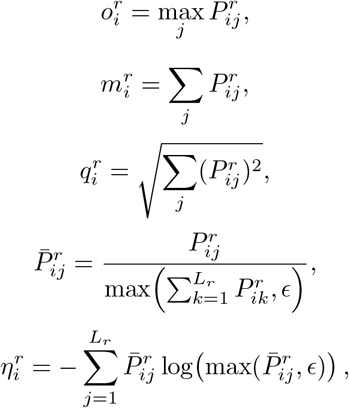

where *ϵ* > 0 ensures numerical stability.

These descriptors summarized maximum pairing confidence, total pairing mass, *l*_2_ mass and entropy. A structure adapter projected them into 32-dimensional embeddings:

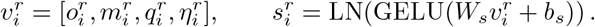

The initial nucleotide representation concatenated sequence, metadata and structural features:

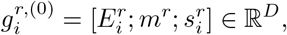

where

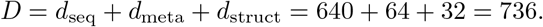

Here, *d*_meta_ and *d*_struct_ denote the metadata and structural embedding dimensions, respectively.

#### Opening and pairwise hybridization branches

The opening branch transformed each structural embedding into a non-negative accessibility-related score,

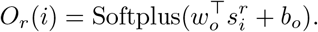

An auxiliary loss encouraged agreement with the RibonanzaNet-SS-derived maximum pairing-confidence feature. The scores entered pairwise logits through learned coefficients *w*_*A*_ and *w*_*B*_, initialized at 1.0 without positivity constraints.

We indexed the *K* = 3 global cross-attention layers by *l* = 0, …, *K* − 1, with 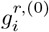 denoting the initial fused representation.At each layer, a pairwise MLP estimated context-dependent hybridization gain:

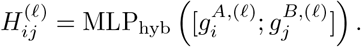

The MLP used a GELU hidden layer and was shared across the global interaction layers. Hybridization gains were recomputed at each layer from its current input representations.

#### Accessibility-aware axial cross-attention

At global cross-attention layer *l* contextual affinity between RNA-A nucleotide *i* and RNA-B nucleotide *j* was calculated as

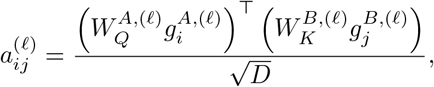

where 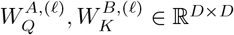 are learnable query and key projections. We z-score-normalized the affinity and hybridization-gain maps within each sample, obtaining 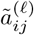 and 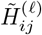The reported configuration combined these terms as

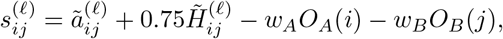

where w_*A*_ and w_*B*_ are learned scalar coefficients. Contextual affinity and hybridization gain had fixed weights of 1.0 and 0.75, and the canonical-pair prior *M*_pair_ was disabled.

The shared score map was normalized separately in the two directions,

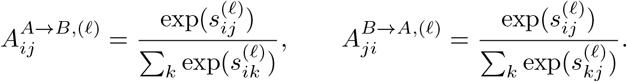

Attention-weighted value projections updated both RNA streams through residual addition and layer normalization. For each query nucleotide, directional attention depended on the opening-related scores of candidate partner positions. The query’s own opening term was constant across those positions and therefore canceled during softmax normalization. The unnormalized pairwise score map retained opening contributions from both RNAs. Three stacked global interaction layers recomputed pairwise logits from the updated representations.

Local inputs used the same sequence, metadata and structural encoders. Their contextual affinity was calculated directly from the fused local representations by a scaled dot product. Pairwise gains and opening-related terms were combined using the same score-fusion rule, with padded positions masked. One bidirectional attention update produced local representations. A learned scale controlled both local-logit injection into the final global map and the local contribution to the pooled pair representation.

#### Pair-level classifier and model outputs

Attention pooling summarized the final global representations of both RNAs. A shared pooling network assigned a score to each nucleotide:

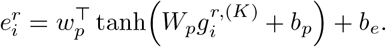

The scores were normalized across nucleotide positions:

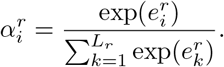

The pooled representation was

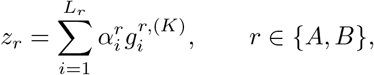

where *W*_*p*_, *w*_*p*_, *b*_*p*_ and b_*e*_ are learnable pooling parameters. When local inputs were available, the joint representation was

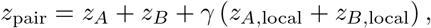

where *γ* was the learned local-window scale. A two-layer MLP with a 128-dimensional GELU hidden layer produced the interaction probability,

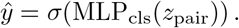

Let *S*^final^ denote the last global score map, s^(*K*−1)^, after adding the scaled local logits at their corresponding global-window coordinates. The model returned 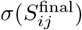 as a nucleotide-pair evidence map. The same learned scale *γ* controlled local-logit injection and the local contribution to the pooled pair representation. Interpretation exports included directional attention, hybridization gains, fused logits, opening scores, structural features and input coordinates.

#### Training objective

The primary objective was binary cross-entropy on RNA-pair interaction labels. An auxiliary loss, weighted by *β*_open_ = 0.001, minimized the mean-squared error between opening scores and detached RibonanzaNet-SS occupancy targets. RibonanzaNet-SS remained frozen in the reported RNA-FM fine-tuning configuration and supplied both structural features and consistency targets. The checkpoint with the highest validation ROC–AUC was selected for testing.

### 4.3 Assessing MARRI hotspot faithfulness

#### 4.3.1 Selection of localized high-confidence hotspots

We selected positive test-set pairs scored by the fixed seed-42 full-model checkpoint. Eligible pairs had predicted probabilities of at least 0.80 and local segments of at least 20 nt on both RNAs. Within each local map, qualifying cells ranked simultaneously in the top 0.5% of fused logits and top 1% of reciprocal attention. Fused logits integrated contextual affinity, hybridization gain and opening penalties. Reciprocal attention was the geometric mean of the two directional attention weights at corresponding nucleotide pairs:

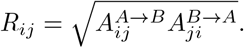

Qualifying cells were grouped using 8-neighbor connectivity, including shared edges and corners. Components required at least three cells. When multiple components qualified, we retained the one with the largest summed standardized fused-logit score. This procedure identified 182 localized high-confidence candidates.

#### 4.3.2 Localized perturbation design and statistical analysis

For each selected pair, we retained the wild-type (WT) input and generated a hotspot-disruption variant. At least three RNA-A nucleotides were changed within a 7-nt neighborhood centered on the hotspot, with each replacement differing from the original base. Where an auxiliary AF3 model provided a local inter-RNA contact, its RNA-A positions guided substitution placement ^55^. AF3 modeling and contact extraction are described in Supplementary Section S.6.

Each pair received two to five matched random-site controls outside the hotspot interval. Controls preserved the number and relative arrangement of substitutions. Random intervals were matched as closely as possible for GC content and dinucleotide composition. All inputs were rescored with the same fixed checkpoint.

For pair *n*, the hotspot effect was

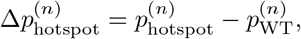

and the matched-control effect was

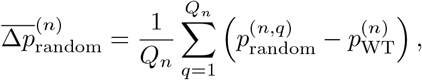

where *Q*_*n*_ is the number of valid random controls for that pair. A one-sided paired Wilcoxon signed-rank test assessed whether hotspot effects were more negative than mean matched-control effects across the 182 pairs. This comparison tested whether the nominated region influenced model scores more strongly than matched changes elsewhere in the input.

### 4.4 Constructing disease-associated candidate RNA–RNA networks

#### 4.4.1. Candidate-pair preparation and window-level MARRI inference

All disease analyses used a fixed MARRI checkpoint and disease-specific candidate pools. Reverse-complement k-mer matches and local-complementarity scores prioritized candidate window combinations. RIblast located local interaction sites within these combinations ^56^. Each retained site was centered in a 256-nt RNA-A/RNA-B window pair, with local intervals stored as relative start and end coordinates.

The training N-clean rule was applied before structural prediction and interaction scoring. Both context windows had to contain exclusively A, C, G and U; ambiguous bases, including boundary-padding N bases, caused exclusion. MARRI then assigned an interaction probability to each retained window pair.

#### 4.4.2 Disease-specific expression backgrounds and candidate sets

##### Diabetic nephropathy

We obtained renal-biopsy expression data from GSE142025 and assigned samples to Normal, Early DN and Advanced DN using the accompanying metadata ^34^. The processed matrix contained quantile-normalized, log_2_-transformed expression values. Stage-specific expressed RNAs required values above 1 in at least 20% of samples. GENCODE supplied RNA-biotype and transcript-region annotations.

The evidence-supported A-class set retained human RNA–RNA edges recorded in RNAInter or NPInter when both partners were expressed in the relevant stage ^35,36^.

The exploratory B-class set used a separate panel of 300 expressed RNAs for each stage. The Normal panel included the 200 most abundant RNAs and 100 RNAs with pronounced DN-associated changes that remained expressed in Normal samples. Each DN panel included 150 highly expressed RNAs, 75 upregulated RNAs and 75 downregulated RNAs. Duplicates were removed and remaining panel positions were filled by expression rank. All unordered non-self combinations yielded 44,850 candidate pairs per stage.

For both classes, RIblast evaluated the 50 highest-priority window combinations per RNA pair. A-class pairs retained up to five highest-ranking sites. B-class pairs retained the best site and up to two additional sites with local-complementarity scores at least 90% of the best-site score.

##### Atherosclerosis

The discovery dataset GSE233402 contained EFA (n = 4), LFA (n = 4) and TCFA (n = 5) carotid plaques ^43^. Genes required counts per million above 1 in at least two samples of one or more groups, retaining 24,511 genes after expression filtering. Of these, 24,488 had estimable differential-expression P values in the TCFA–LFA comparison. PyDESeq2 fitted a plaque-phenotype design, with TCFA versus LFA as the primary contrast ^57^. Stringent differential expression required Benjamini–Hochberg FDR < 0.05 and | log_2_ FC| > 1. Reactome enrichment of TCFA-upregulated genes used the 24,488 genes with estimable P values as the background, with g:Profiler multiple-testing correction ^58,59^. Prespecified macrophage, smooth-muscle and antigen-presentation modules were summarized by sample-level mean expression z scores.

GSE120521 supplied stable and unstable plaque regions from four patients for an external expression comparison ^46^. We calculated paired patient-level differences in log_2_(FPKM + 1). Stable Ensembl identifiers were matched after excluding 45 identifiers with X/Y pseudoautosomal ambiguity. Directional agreement compared discovery TCFA–LFA fold changes with unstable–stable effects.

We selected one common panel of 300 expressed RNAs from pooled abundance across the 13 discovery samples, including 150 protein-coding RNAs and 150 lncRNAs. All three plaque groups therefore shared candidate RNAs, local windows and fixed MARRI scores. We considered all 44,850 unordered non-self pairs. Local-complementarity prioritization and RIblast localization generated 256-nt windows, which underwent N-clean filtering before inference.

#### 4.4.3 RNA-pair scoring and expression-conditioned network analysis

Each scorable RNA pair received the maximum probability across its retained windows:

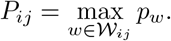

Here, *W*_*ij*_ is the set of retained scorable windows and *p*_*w*_ is the window-level prediction. The score therefore represented the strongest model support among the prioritized sites for that pair.

For DN B-class networks, stage-specific availability was calculated from mean expression:

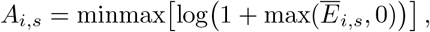

Where *Ē*_*i,s*_ is RNA *i’*s mean normalized expression in stage s. Min–max scaling was performed separately within each stage over the union of B-class RNAs. Edge weights were

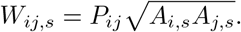

Weights from all three stages defined a pooled 95th-percentile threshold of *q*_95_ = 0.635859. Stage-specific backbones retained edges meeting or exceeding this shared threshold.

DN networks were displayed using force-directed layouts. Node color encoded RNA type, size encoded degree and opacity encoded stage-specific availability. Weighted modules were identified with Louvain community detection. Within-module connectivity and participation were calculated using weighted connectivity ^60^:

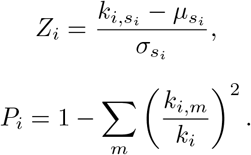

Here, 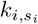 is RNA *i’*s edge-weight sum within its assigned module, and 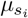 and 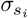 describe that module’s distribution. In the participation coefficient, *k*_*i,m*_ is connectivity to module *m* and *k*_*i*_ is total weighted connectivity. Our four-role grouping used boundaries *Z*_*i*_ = 2.5 and *P*_*i*_ = 0.62. Network hubs exceeded both boundaries; connectors met the participation boundary but had *Z*_*i*_ < 2.5. Module hubs had *Z*_*i*_ ≥ 2.5 and *P*_*i*_ < 0.62; remaining nodes were peripheral.

For functional analysis, we selected the top-5% weighted-degree RNAs from each stage-specific A-class network. Reactome over-representation analysis used the corresponding network RNA set as a custom background and g:Profiler FDR correction ^58,59^. The displayed pathways met adjusted *P* < 0.05.

For atherosclerosis, RNA availability was calculated from sample-level log(1 + CPM), scaled against the maximum over the common panel and all discovery samples. Sample-level weights were

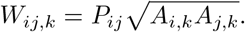

Group weights were the sample means within each plaque phenotype. A pooled 95th-percentile threshold, *q*_95_ = 0.499897, defined the shared backbones. Sensitivity analyses used pooled *q*_90_ and *q*_97.5_ thresholds. Networks shared coordinates; node size denoted degree and opacity denoted phenotype-level mean availability.

TCFA–LFA edge differences were tested using exact phenotype-label permutations, followed by Benjamini–Hochberg correction across scored pairs. We used 2,000 bootstrap resamples for pointwise confidence intervals and leave-one-out direction stability to prioritize effects. The resulting comparisons combined fixed RNA-pair scores with sample-or group-specific expression availability.

### 4.5 Training protocol and model outputs

Unless otherwise specified, data were partitioned into training, validation and test sets at an 8:1:1 ratio. The training dataset was approximately balanced by dynamically sampling negative pairs in each epoch, while preserving the PN:NP:NN ratio of 50:25:25. Training supported distributed execution and mixed-precision optimization.

We compared four backbone configurations by independently freezing or fine-tuning RNA-FM and RibonanzaNet-SS, under both static and dynamic negative sampling (Supplementary Table S2). Based on these experiments, we adopted RNA-FM fine-tuning with frozen RibonanzaNet-SS and dynamic negative sampling as the main configuration. This configuration achieved the highest mean ROC–AUC among those evaluated.

Models were trained for up to 100 epochs, with a minimum training duration of 10 epochs. Validation ROC–AUC was evaluated after each epoch, and any new highest value updated the saved checkpoint. The early-stopping counter was reset when validation ROC–AUC improved by at least 0.005 over the reference value recorded at its previous reset. Otherwise, the counter increased, and training stopped after five consecutive non-reset rounds once the minimum training duration was reached. Thus, smaller improvements could update the best checkpoint without resetting patience. The checkpoint with the highest validation ROC–AUC was reloaded for test evaluation.

At inference, MARRI returned RNA-pair interaction probabilities and the interpretation outputs described in Section 4.2, including directional attention maps, hybridization gains, fused logits, opening scores and structural features.

## 5 Data and code availability

The source code and datasets will be made publicly available upon publication.

## 6 Acknowledgments

B.W. conducted part of this work as a remote summer research intern at Brigham and Women’s Hospital and Harvard Medical School. Z.M. conducted part of this work as a visiting PhD researcher at Brigham and Women’s Hospital and Harvard Medical School. This work was supported in part by AHA MERIT grant 1185447; by EU Horizon Health 2021 grant 101057619; and by a Ruggeri Family Network Medicine grant to J.L.

## 7 Author contributions

B.W.: Conceptualization, Investigation, Methodology, Data Curation, Software, Resources, Formal Analysis, Visualization, Validation, Writing Manuscript; H.L.: Methodology, Resources, Manuscript Proofreading; Z.M.: Resources, Manuscript Proofreading; G.C.: Data Curation, Manuscript Proofreading; X.W.: Manuscript Proofreading; Y.C.: Resources, Manuscript Proofreading; J.S.: Conceptualization, Resources, Manuscript Proofreading; J.L. & H.X.: Conceptualization, Investigation, Methodology, Software, Resources, Supervision, Project Administration, Revising Manuscript. All authors read and approved the final manuscript.

## Materials & correspondence

Correspondence should be addressed to. Requests for materials should be addressed to Bo Wang.

