## supplements for "MARRI: Mechanism-Aware Learning of RNA–RNA Interactions with Foundation Models"

### Supplementary Material

#### Contents

S.1 Detailed architecture of MARRI **S2**

S.2 Computational resources **S2**

S.3 Training across random seeds **S3**

S.4 Generalization to unseen RNAs and interaction regions **S4**

S.5 Discrimination against challenging negatives **S5**

S.6 Structural information for hotspot analysis **S5**

S.7 RIME performance across random seeds **S5**

#### S.1 Detailed architecture of MARRI

MARRI applies a shared encoding procedure to both RNA partners, combining sequence representations, structural summaries and categorical annotations (Supplementary Fig. S1). Global inputs retain flanking context; local inputs represent the selected candidate interaction regions. RNA/gene-type categories are protein-coding, lncRNA, miRNA, ncRNA, pseudogene, snRNA, rRNA, snoRNA, tRNA, other and unknown. Transcript-region categories are exon, 5′ UTR, coding sequence, 3′ UTR, intron and unknown. The annotation vocabularies are predefined, and embedding parameters are learned during training.

Frozen RibonanzaNet-SS generates intramolecular pair-confidence matrices. Per-nucleotide occupancy, total pairing mass, $\ell_2$ mass and entropy are projected into structural embeddings, from which an opening branch learns non-negative accessibility-related scores. At each cross-attention layer, a pairwise MLP recomputes hybridization gains from the current RNA representations. Contextual affinity, hybridization gain and opening penalties contribute to pairwise scoring as both RNA streams are updated. Attention pooling and an MLP classifier produce the RNA-pair interaction probability.

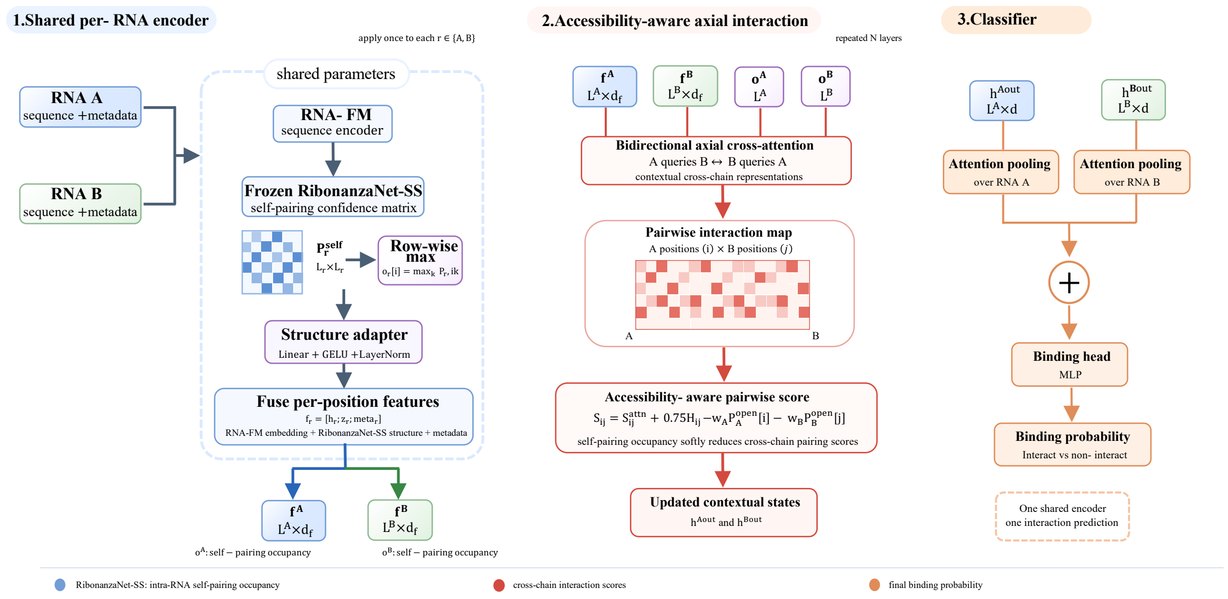


**Figure S1: Detailed architecture of MARRI.** **1**, shared per-RNA encoding. RNA-FM generates global and local representations. Frozen RibonanzaNet-SS provides intramolecular pair-confidence matrices, summarized into nucleotide-level structural features and projected by a structure adapter. Sequence, structural and metadata features form per-position representations. **2**, accessibility-aware axial interaction. Bidirectional cross-attention combines contextual affinity, hybridization gain and opening penalties. Stacked layers update both RNA representations and recompute pairwise evidence maps. **3**, classification. Attention pooling aggregates RNA representations, which are combined and passed to an MLP classifier to obtain the interaction probability.

#### S.2 Computational resources

The highest-performing configuration fine-tuned RNA-FM, froze RibonanzaNet-SS and used dynamic negative resampling. Training used four NVIDIA H100 80GB GPUs with distributed data parallelism, one process per GPU, bf16 precision and batch size 4 per GPU (global batch size 16). Training logs provided the timings in Supplementary Table S1.

An epoch comprised 8,029 optimization steps over 128,452 training pairs and took an average of 53.9 min (2.92 steps/s; 0.342 s/step across four GPUs). Validation over 16,056 pairs, sharded across GPUs, took approximately 5.3 min. The highest validation ROC–AUC occurred at epoch 23, after 22.75 h; training stopped at epoch 24 after 23.74 h. Checkpoint selection and early-stopping-counter updates followed the separate rules described in Methods. Test inference and interpretation export for 16,058 pairs took 35.5 min on one GPU at batch size 4 (1.89 batches/s). Exports included attention maps, hybridization gains and opening penalties.

**Table S1:** Computational resources for training and evaluating the best-performing model (RNA-FM fine-tuned, RibonanzaNet-SS frozen, dynamic negative resampling).

| Item | Value |
|---|---|
| Hardware | 4× NVIDIA H100 80GB HBM3 (DDP, 1 process/GPU) |
| Precision | bf16 |
| Batch size | 4/GPU (global 16) |
| Training throughput | 2.92 it/s (0.342 s/step) |
| Time per training epoch | 53.9 min (8,029 steps) |
| Time per validation pass | 5.3 min (16,056 pairs, sharded) |
| Epochs to best checkpoint | 23 |
| Cumulative training time to best checkpoint | 22.75 h |
| Total training time (incl. early-stop patience) | 23.74 h |
| Test + interpretability export (1 GPU) | 35.5 min (16,058 pairs, 1.89 it/s) |

#### S.3 Training across random seeds

We trained the eight backbone/resampling configurations and four component-ablation configurations using seeds 42, 43 and 44. Tables 2 and S2 report mean test ROC–AUC and across-seed standard deviation. Tables S3 and S4 provide individual values and validation-selected epochs.

All runs used batch size 4 per GPU across four GPUs (global batch size 16), learning rate $1 \times 10^{-4}$, AdamW with weight decay $1 \times 10^{-4}$ and bf16 precision. Training continued for at least 10 epochs, with early-stopping patience 5 and an AUC-reference improvement threshold of 0.005. Checkpoints were saved at the highest validation ROC–AUC, independently of the early-stopping counter, as described in Methods. Within each configuration, only the random seed was varied; the code and hyperparameters were held constant, while data partitions were fixed within each run but varied across seeds. Seeds governed weight initialization, data shuffling and dynamic negative sampling where enabled.

Dynamic sampling redrew training negatives each epoch at PN:NP:NN $= 50 : 25 : 25$; static sampling retained one negative set throughout training. Dynamic sampling improved mean ROC–AUC in all four backbone configurations. Gains were 0.89 percentage points with both backbones frozen (1.2% relative), 0.77 with RNA-FM fine-tuning (1.1%), 0.66 with RibonanzaNet-SS fine-tuning (0.9%), and 0.46 with both fine-tuned (0.6%).

**Table S2:** Effect of epoch-wise dynamic negative resampling relative to a static (fixed) negative training set, across the four backbone fine-tuning configurations. Values are percentages. Static and Dynamic are mean ± standard deviation of test ROC–AUC over three random seeds (42, 43, 44); Δ is the change in the means, in percentage points.

| Backbone configuration | Static | Dynamic | Δ ROC–AUC |
|---|---|---|---|
| **Both frozen** | 72.03 ± 0.45 | 72.92 ± 0.44 | +0.89 |
| RibonanzaNet-SS fine-tuned | 71.72 ± 0.52 | 72.39 ± 1.20 | +0.66 |
| RNA-FM fine-tuned | 73.50 ± 0.63 | 74.27 ± 0.95 | +0.77 |
| Both fine-tuned | 72.91 ± 0.22 | 73.37 ± 0.91 | +0.46 |

**Table S3:** Per-seed test ROC–AUC for the eight main-matrix (backbone freeze/fine-tune) configurations, underlying the mean ± SD values in Table S2. Values are percentages. Rows are grouped by backbone configuration, static then dynamic. "Best epoch" is the epoch at which the validation-ROC–AUC-selected checkpoint occurred for that seed (early stopping: minimum 10 epochs, patience 5 epochs, $\Delta \geq 0.005$).

| | Test ROC–AUC | | | | Best epoch | | |
|---|---|---|---|---|---|---|---|
| Configuration | Seed 42 | Seed 43 | Seed 44 | Mean ± SD | 42 | 43 | 44 |
| Both frozen, static | 72.41 | 72.15 | 71.54 | 72.03 ± 0.45 | 8 | 10 | 9 |
| Both frozen, dynamic | 73.40 | 72.52 | 72.84 | 72.92 ± 0.44 | 13 | 10 | 15 |
| RibonanzaNet-SS fine-tuned, static | 72.08 | 71.97 | 71.12 | 71.72 ± 0.52 | 13 | 11 | 12 |
| RibonanzaNet-SS fine-tuned, dynamic | 73.20 | 72.95 | 71.01 | 72.39 ± 1.20 | 13 | 15 | 11 |
| RNA-FM fine-tuned, static | 74.12 | 73.52 | 72.85 | 73.50 ± 0.63 | 23 | 12 | 13 |
| RNA-FM fine-tuned, dynamic (full model) | **75.30** | 74.09 | 73.42 | **74.27 ± 0.95** | 23 | 15 | 18 |
| Both fine-tuned, static | 73.16 | 72.82 | 72.75 | 72.91 ± 0.22 | 12 | 10 | 12 |
| Both fine-tuned, dynamic | 74.34 | 73.23 | 72.53 | 73.37 ± 0.91 | 17 | 11 | 11 |

**Table S4:** Per-seed test ROC–AUC for the full model and its four component ablations, underlying the mean ± SD values in Table 2. Values are percentages. Row order matches Table 2. "Best epoch" is the epoch at which the validation-ROC–AUC-selected checkpoint occurred for that seed (early stopping: minimum 10 epochs, patience 5 epochs, $\Delta \geq 0.005$).

| | Test ROC–AUC | | | | Best epoch | | |
|---|---|---|---|---|---|---|---|
| Configuration | Seed 42 | Seed 43 | Seed 44 | Mean ± SD | 42 | 43 | 44 |
| Full model | **75.30** | 74.09 | 73.42 | **74.27 ± 0.95** | 23 | 15 | 18 |
| w/o local interaction window | 70.27 | 70.20 | 69.45 | 69.97 ± 0.46 | 20 | 23 | 19 |
| w/o metadata embedding | 72.49 | 72.27 | 71.30 | 72.02 ± 0.63 | 18 | 18 | 18 |
| w/o RibonanzaNet-SS structural features | 73.82 | 73.23 | 73.46 | 73.50 ± 0.30 | 15 | 9 | 18 |
| w/o hybridization gain | 74.13 | 73.29 | 73.72 | 73.72 ± 0.42 | 13 | 15 | 15 |

#### S.4 Generalization to unseen RNAs and interaction regions

We evaluated MARRI using three non-random held-out strategies defined at the gene, RNA-pair and interaction-locus levels. A model was independently retrained for each setting using the same backbone configuration and negative-resampling protocol. Supplementary Table S5 reports the complete results across all six evaluation metrics.

**Table S5:** Generalization under non-random held-out splits. Each model was retrained on its corresponding split with seed 42, RNA-FM fine-tuning, frozen RibonanzaNet-SS and dynamic negative resampling. Values are percentages.

| Split strategy | Generalization tested | ROC–AUC | PR–AUC | Accuracy | Precision | Recall | F1-score |
|---|---|---|---|---|---|---|---|
| Gene-held-out | Unseen RNA/gene | 66.30 | 17.50 | 56.56 | 13.85 | 68.01 | 23.01 |
| Interaction-pair-held-out | Unseen RNA–RNA pair | 74.22 | 32.23 | 63.74 | 23.55 | 72.09 | 35.50 |
| Interaction-locus-held-out | Unseen local interaction locus | 75.02 | 33.51 | 62.99 | 23.56 | 74.58 | 35.81 |

#### S.5 Discrimination against challenging negatives

We compared positive interactions with partner-swapped negatives (PN), alternative local regions from the same RNA pairs (NP) and random background pairs (NN). These categories test discrimination against alternatives that retain different amounts of interaction-associated context.

ROC–AUC was highest against NN (0.8730), followed by NP (0.7733) and PN (0.6529; Supplementary Table S6). Partner-swapped negatives therefore posed the greatest challenge, showing that performance against random pairs alone does not capture discrimination among plausible partners.

At a common threshold of 0.5, F1-scores ranged from 0.7062 to 0.7913. Comparison-specific threshold optimization increased F1 to 0.8074, 0.9034 and 0.9164 for PN, NP and NN, respectively. These analyses illustrate how an operating threshold changes the precision–recall balance within a defined candidate set. In applications, a threshold could be chosen on separate validation data to favor sensitivity or precision according to screening needs. Here, thresholds were optimized on the evaluated data, so the resulting F1 values describe post hoc operating points.

**Table S6:** Discrimination of positive pairs against PN, NP and NN negatives. AP denotes average precision. Default F1 uses threshold 0.5; best F1 and its threshold were selected separately within each evaluated comparison.

| Comparison | ROC–AUC | AP | F1 ($\tau_0$=0.5) | Best F1 | Best threshold |
|---|---|---|---|---|---|
| PP vs PN | 0.6529 | 0.7833 | 0.7062 | 0.8074 | 0.1689 |
| PP vs NP | 0.7733 | 0.9164 | 0.7733 | 0.9034 | 0.1572 |
| PP vs NN | 0.8730 | 0.9605 | 0.7913 | 0.9164 | 0.1934 |

#### S.6 Structural information for hotspot analysis

AlphaFold3 (AF3)[55] was used to assist substitution placement in hotspot-disruption variants. Inter-RNA base contacts were extracted from top-ranked AF3 models using a 4.5 Å base-heavy-atom distance cutoff.

For each selected pair, we retained the wild-type (WT) input and generated a hotspot-disruption variant. At least three RNA-A nucleotides were changed within a 7-nt neighborhood centered on the hotspot, with each replacement differing from the original base. Where an auxiliary AF3 model provided a local inter-RNA contact, its RNA-A positions guided substitution placement.

#### S.7 RIME performance across random seeds

Supplementary Table S7 reports RIME test performance for seeds 42, 43 and 44. Aggregate performance is reported in the main text.

**Table S7:** RIME test performance for individual random seeds. Values are percentages.

| Seed | ROC–AUC (%) | PR–AUC (%) | Accuracy (%) | Precision (%) | Recall (%) | F1-score (%) |
|---|---|---|---|---|---|---|
| 42 | 61.99 | 62.00 | 57.97 | 57.11 | 62.01 | 59.46 |
| 43 | 62.49 | 62.41 | 58.55 | 58.01 | 62.10 | 59.99 |
| 44 | 61.75 | 61.56 | 57.63 | 56.64 | 61.20 | 58.83 |
